# A genome-wide CRISPR activation map of surface protein expression in human CD4 T cells

**DOI:** 10.64898/2026.08.08.743670

**Authors:** Yutong V. Wang, Junha Park, Min Cheol Kim, Thomas Mazumder, Krishna Sonpal, Matin Bikaran, Zachary Steinhart, Ralf Schmidt, Yang Sun, Su-Hyeon Lee, Alexander Marson, Jimmie Ye, Byungjin Hwang

## Abstract

Surface proteins define T cell identity and function, but the abundance of each protein is not determined by transcription alone. Existing genome-wide CRISPR screens in primary human T cells either profile the transcriptome or isolate cells based on a single functional or protein phenotype. Here we present SCITO-Perturb-seq, a novel platform that couples combinatorial-indexed single-cell cytometry sequencing with pooled CRISPR activation (CRISPRa) to map the causal regulation of 201 surface proteins across 3.6 million human CD4 T cells. We find that 16% of activated genes significantly alter the expression of at least one surface protein. By applying semi-nonnegative matrix factorization to the perturbation effect matrix, we identified five modules corresponding to known CD4 T cell states. Notably, these modules group surface proteins by their shared response to perturbation, revealing coordinated regulation of proteins that are not co-expressed in unperturbed cells. SCITO-Perturb-seq represents the first genome-wide CRISPRa screen paired with direct, high-dimensional surface protein profiling, providing a comprehensive regulatory map of the CD4 T cell surface proteome.

## Introduction

Surface proteins on T cells mediate diverse functions, including antigen recognition^1^, signal transduction^2^, physical adhesion^3^, metabolic transport^4^, and environmental sensing^5^. They are the primary markers used to define T cell lineage, differentiation outcomes^6^, and functional states^7^. While a surface protein’s abundance begins with transcription, it is ultimately shaped by post-transcriptional and post-translational mechanisms, making transcript levels an incomplete proxy for actual cell-surface expression. Furthermore, cell states are inherently defined by combinations of multiple proteins rather than any single marker. For instance, PD-1 marks both recent activation and chronic exhaustion, and which state it indicates depends on other surface proteins a cell co-expresses^8^. To causally understand how the genome shapes these multi-protein cell states, we require a functional genomic platform that couples genome-wide CRISPR perturbations with highly multiplexed, single-cell protein profiling.

Pooled CRISPR knockout, activation, and inhibition screens in primary T cells have reached genome scale^9,10^, but these approaches remain unable to map high-dimensional surface protein profiles, capturing instead either a single functional trait or the high-dimensional transcriptome. Functional screens either select cells based on survival phenotypes such as proliferation^9^, persistence^11^, or effector function^12^, or by individual protein expression as proxies for function^13^, before using bulk sequencing to quantify sgRNA enrichment. A 6,000-sgRNA library against approximately 1,350 transcription factors and chromatin modifiers mapped regulators of IL2RA^14^, and a genome-wide knockout screen mapped regulators of FOXP3 by sorting primary human CD4 T cells on intracellular FOXP3 protein^15^. Because these screens pool sorted cells for bulk sequencing, they recover only the sgRNA frequency in each bin and cannot link genotype to phenotype within individual cells. Perturb-seq^16,17^ overcomes this by integrating CRISPR perturbation with single-cell RNA sequencing, preserving the direct coupling between a genetic perturbation and the expression profile of the perturbed cell. While recent work has demonstrated the feasibility of genome-wide Perturb-seq in primary human CD4+ T cells^18^ and cell lines^19,20,21^, these transcriptomic approaches cannot account for the post-translational processes that shape the surface proteome-the layer where T cell identity and functional responses are executed.

Technologies capable of pairing single-cell CRISPR perturbations with multiplexed surface protein profiling do exist, but none have successfully scaled to genome-wide libraries. For instance, multimodal approaches like ECCITE-seq^22,23^ and Perturb-CITE-seq^24^ couple CRISPR perturbations with joint RNA and surface protein measurements within individual cells. Perturb-CITE-seq profiled the effects of knocking out 248 genes on 20 surface proteins in melanoma-TIL co-cultures^24^. More recently, Perturb-icCITE-seq advanced multiplexed single-cell profiling by coupling CRISPR perturbations of 296 targeted candidate genes with the measurement of over 300 surface and intracellular proteins during primary human CD4 T cell differentiation^15^. However, scaling these methods to a genome-wide library is impractical due to the cell throughput constraints of commercial microfluidic platforms and the high per-cell cost of transcriptome sequencing. Because a genome-scale screen requires profiling millions of cells to ensure sufficient coverage per sgRNA, the combination of physical cell-recovery limits and deep sequencing overhead makes sample preparation and processing financially and logistically prohibitive. Alternatively, mass cytometry approaches like Pro-Code^25^ assign sgRNAs to combinatorial protein barcodes that can be decoded by cyTOF but are physically limited by up to 50 detection channels. This structural constraint limits these platforms to profiling tens of surface proteins across hundreds of perturbations, preventing them from scaling to genome-wide libraries. To map the genetic determinants of surface protein expression at scale, a new framework is required that overcomes the throughput limitations of commercial microfluidics and the multiplexing constraints of mass spectrometry.

Here, we present SCITO-Perturb-seq, a platform that overcomes existing throughput and targeting constraints by combining pooled CRISPR activation (CRISPRa) with combinatorial-indexed single-cell cytometry^26^. This approach utilizes commercially available microfluidics, but increases cell throughput per channel approximately 20-fold and drastically lowers per-cell sequencing costs relative to transcriptome-wide profiling. Furthermore, while CRISPR knockout (CRISPRn) and inhibition (CRISPRi) screens can fail to resolve regulators of surface proteins with long half-lives due to the persistence of pre-existing proteins, CRISPRa circumvents this by directly testing whether gene activation is sufficient to drive surface expression. Utilizing SCITO-Perturb-seq, we profiled 201 surface proteins in response to the CRISPRa targeting of 18,325 genes across 3.6 million primary human CD4+ T cells. We estimate that 16% of perturbed genes significantly alter at least one surface protein. These regulatory effects were robustly validated by self-targeted concordance, split-half reliability, and cross-screen comparison to published transcriptional and single-protein datasets^14,18^.

Finally, semi-nonnegative matrix factorization of the gene-by-protein effect matrix identified five modules, each pairing a set of gene perturbations with a surface-protein signature that maps to a known T cell phenotype. Ultimately, SCITO-Perturb-seq surmounts long-standing technological bottlenecks to deliver the first genome-wide causal map linking gene activation directly to the surface proteome of primary human T cells.

## RESULTS

### Developing SCITO-Perturb-seq for genome-wide mapping of surface protein expression

We developed SCITO-Perturb-seq to couple genome-wide CRISPR perturbations with SCITO-seq, a droplet-based single-cell cytometry assay that profiles surface proteins in more than 150,000 cells per reaction using combinatorial indexing^26,27^ (**Fig. 1a**). To adapt SCITO-seq’s combinatorial indexing workflow to capture sgRNA identities at a genome-wide scale, we split the sgRNA library into ten non-overlapping pools. We placed the two sgRNAs targeting each gene into different pools, ensuring that every gene was covered across two independent pools and measured by two independent sgRNAs. Accordingly, cells were separately transduced with each sgRNA pool and stained with a 201-antibody panel, where the antibodies for each respective pool carried a unique, pool-specific oligonucleotide barcode embedded in their ADT sequence. We then mixed all ten pools to load them together onto a standard 10X Genomics microfluidic chip for single-cell processing. Within this workflow, each cell is uniquely identified by the combination of its droplet barcode and its pool-specific antibody barcode. This combinatorial index ensures that even when multiple cells occupy the same droplet, they can be uniquely resolved by their different pool barcodes, allowing the linking of each single-cell surface proteome to its specific sgRNA perturbation.

**Fig. 1.**
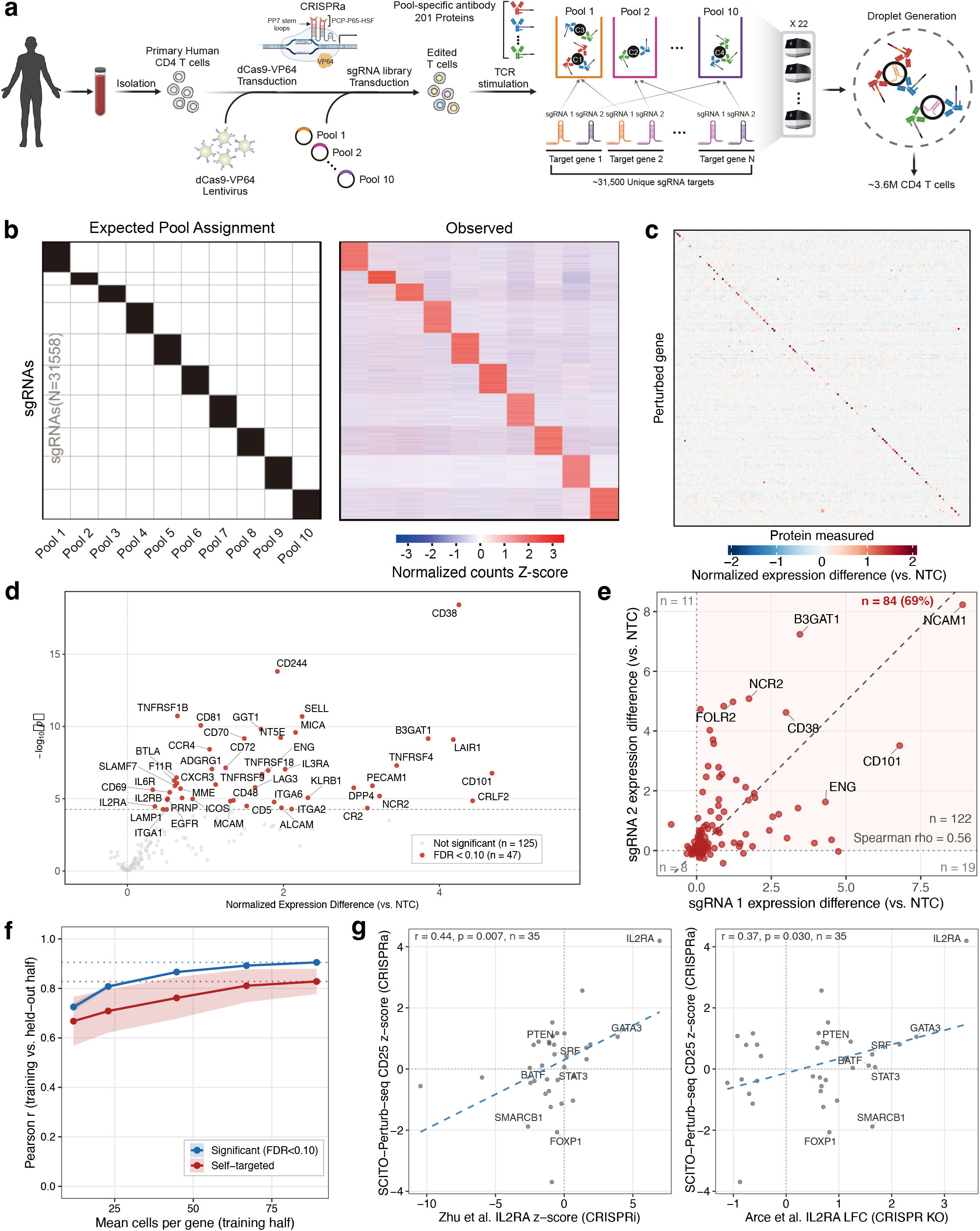
Genome-wide single-cell surface protein screen of primary human CD4 T cells. (a) Experimental design for genome-wide SCITO-Perturb-seq. Primary human CD4 T cells were transduced with a CRISPRa library and profiled for 201 surface proteins across 10 pools, capturing ∼3.6M cells with simultaneous genetic perturbation and high-dimensional surface phenotyping. (b) Expected (left) and observed (right) pool assignment for 31,558 sgRNAs across 10 pools. (c) Heatmap of normalized expression differences (vs. NTC) for 172 self-targeted gene-protein pairs. Rows (perturbed gene) and columns (measured protein) are matched and ordered alphabetically after deduplication to one entry per gene and per protein. Diagonal cells show the self-targeted effect. Off-diagonal cells show effects on non-cognate proteins. (d) Volcano plot of the same 172 self-targeted gene-protein pairs at the gene level. x-axis: normalized expression difference vs. NTC; y-axis: −log₁ ₀ (p-value). Red points (n = 47): per-protein FDR < 0.10; all are labeled by gene name. Dashed horizontal line: FDR = 0.10 threshold. (e) Concordance between the two independent sgRNAs targeting each self-targeted gene. Each point is one self-targeted gene-protein pair, plotted by the normalized expression difference for sgRNA 1 (x-axis) vs. sgRNA 2 (y-axis), for all genes with two evaluable sgRNAs (n = 122 gene-protein pairs, 112 unique genes). Bold red text: both sgRNAs positive (n = 84, 68.9%). Corner annotations show quadrant counts. Dashed line: y = x. Spearman ρ = 0.56. (f) Power analysis for effect-size reliability. Pearson r between normalized expression differences estimated from a downsampled training half and a full-size held-out half, for two subsets of gene-protein pairs: significant pairs (per-protein FDR < 0.10; blue) and self-targeted pairs (red). For each target gene, cells were randomly split 50/50 into training and held-out halves; the training half was progressively downsampled. Lines: means across 10 independent random splits; shaded bands: ±1 s.d. Dotted horizontal lines mark the ceiling reliability at full training-half size (∼89 cells per gene on average). (g) External validation of CD25 (IL2RA) regulators against two published loss-of-function screens. Shown are the 35 genes significant in Arce et al. (CRISPR-KO FACS FDR < 0.05) with data available in all three datasets. y-axis: SCITO-Perturb-seq CRISPRa CD25 Z-score. Left panel x-axis: Zhu et al. CRISPRi IL2RA Z-score in resting T cells (positive = promotes IL2RA; r = 0.44, p = 0.007, n = 35). Right panel x-axis: Arce et al. CRISPR-KO FACS IL2RA LFC at 72-hour stimulation (positive = promotes IL2RA; r = 0.37, p = 0.030, n = 35). Top 8 genes by Arce FDR are labeled.

As a positive control, we activated 22 genes encoding surface proteins with two sgRNAs each, which recovered 29,244 cells at a median of 590 ADT UMI per cell and confirmed that CRISPRa upregulated the targeted proteins (**Supplementary Fig. 1, 2a, b**). After optimizing cell loading based on the pilot experiment (**Supplementary Fig. 2c**), we applied SCITO-Perturb-seq to profile a genome-wide library targeting 18,325 genes with two sgRNAs each. The custom panel measured 201 surface proteins spanning T cell lineage, differentiation, effector function, and exhaustion, plus 7 isotype controls (**Supplementary Table**). We loaded 500,000 cells per channel across 22 channels and recovered approximately 3.6 million primary human CD4 T cells, each expressing at least one of 31,558 detected sgRNAs. This corresponds to approximately 164,000 cells per channel, about 20 times the per-channel yield of standard Chromium. Median recovery was 373,000 cells per pool at 947 ADT UMI per cell. After filtering on ADT counts and excluding one low-quality pool, 3,134,864 cells across nine pools were retained (**Methods**). Applying a minimum threshold of 20 cells per gene, 17,336 genes were retained for analysis, with a median of 150 cells per gene (range 20 to 1,266) evaluated against 26,602 non-targeting control (NTC) cells.

Across all ten pools, 84.6% of sgRNAs were recovered in the pool to which they had been assigned by design (**Fig. 1b**), confirming that the pool-specific barcodes correctly resolved guide identity. To assess biological sensitivity, we examined self-targeted gene-protein pairs, in which CRISPRa activates the gene encoding the measured surface protein (for example, a CD38 sgRNA against CD38 protein). For each pair we computed the normalized expression difference, defined as the mean normalized protein expression in perturbed cells minus that in NTC cells (**Methods**). After cell-count filtering and reduction to one entry per gene and per protein, the self-targeted set comprised 172 gene-protein pairs. These pairs were strongly shifted toward positive values, with 86.0% showing increased expression. The heatmap showed self-targeted diagonal effects elevated above off-diagonal entries (**Fig. 1c**), and 47 of the 172 pairs reached per-protein FDR < 0.10 (**Fig. 1d**). The 125 pairs below this threshold remained predominantly positive (80.8%, binomial p = 9.9 × 10^-13^) and came from genes with fewer assigned cells than the significant pairs (median 130 versus 234; Mann-Whitney p = 1.5 × 10^-8^; **Supplementary Fig. 3**), consistent with limited power rather than absent effects. To assess concordance between the two independent sgRNAs targeting each self-targeted gene, we compared their effects on the cognate protein for all genes with both guides evaluable (n = 122 pairs, 112 genes). Both guides produced positive effects for 68.9% of pairs (n = 84/122; Spearman ρ = 0.56; **Fig. 1e**). Most discordant cases reflected one responsive and one non-responsive guide rather than failure of both.

To assess whether effect-size estimates are reproducible at current cell counts, we performed a split-half reliability analysis. For each gene, we randomly partitioned cells into a training half and a held-out half, computed normalized expression differences independently in each, and downsampled the training half to trace reproducibility against cell number (**Methods**). Self-targeted pairs provide an unbiased estimate because they are selected by gene identity, not by observed significance. They reached Pearson r = 0.83 between halves at full training size (∼89 cells per gene; **Fig. 1f**). Pairs reaching per-protein FDR < 0.10 reached r = 0.91 and stayed above 0.87 when the training half was halved to ∼45 cells. For each gene-protein pair we summarized the effect as a perturbation Z-score, the normalized expression difference divided by its Welch standard error (**Methods**). To validate effect estimates against external data, we compared SCITO-Perturb-seq CD25 (IL2RA) Z-scores with two published loss-of-function screens. The Zhu et al. CRISPRi screen^18^ tracks IL2RA expression at single-cell resolution, whereas the Arce et al. CRISPR-KO screen^14^ relies on phenotypic selection, sorting cells by FACS based on IL2RA surface protein expression to score guide enrichment across bulk sort bins. We restricted the comparison to genes significant in the Arce screen and present in all three datasets, which defines a high-confidence set of IL2RA regulators. With all three screens oriented so that positive values mark positive regulators of IL2RA, CRISPRa effects on CD25 correlated with loss-of-function effects on IL2RA in both Zhu et al. CRISPRi (resting T cells, r = 0.44, p = 0.007) and Arce et al. CRISPR-KO FACS (72-hour stimulation, r = 0.37, p = 0.030; **Fig. 1g**).

### Genome-wide perturbation map of 201 surface proteins in CD4+ T cells

Before examining perturbation effects, we characterized how the 201 surface proteins co-vary across unperturbed cells. We computed pairwise Pearson correlations among all 201 proteins across 26,602 NTC cells and clustered them hierarchically into five groups (**Fig. 2a, left; Supplementary Table**). We labeled the groups by their dominant membership as Core (n = 24, e.g., CD4, CD45), Activation (n = 42, e.g., CD25, CD71), Adhesion (n = 39, e.g., CD62L, CD31), Innate-like (n = 57, e.g., CD56, CD16, CD161), and Other (n = 39), a functionally heterogeneous group with no dominant annotation. Within-group correlations exceeded between-group correlations in every group, strongest in Core and weakest in Innate-like. These groups form a baseline reference, defined without perturbation information. Of 17,336 genes testable after cell-count filtering (≥20 cells per gene; **Methods**), 2,848 (16%) significantly altered the expression of at least one surface protein when activated (per-protein FDR < 0.10). When the perturbation Z-score matrix was ordered by the baseline NTC protein groups (**Fig. 2a, right**), proteins in the same baseline group displayed highly similar perturbation responses, indicating that baseline co-expression is associated with shared regulation.

**Fig 2.**
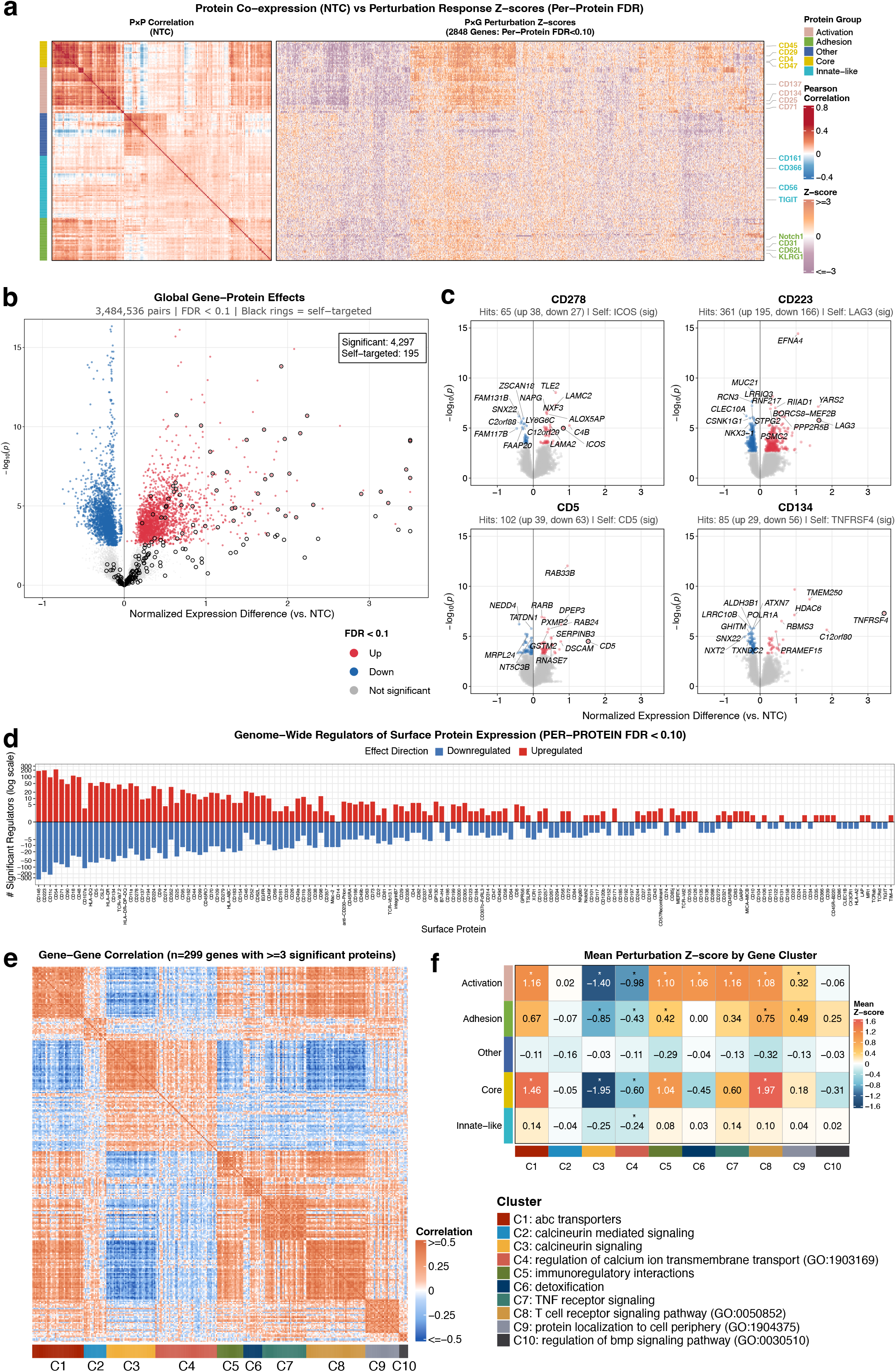
Perturbation mapping reveals pathway-specific control of the CD4 T cell surface proteome. (a) Left: protein-protein correlation from NTC baseline co-expression, with hierarchical clustering (Ward’s D2) defining five protein groups. Right: perturbation Z-score matrix for 2,848 genes with at least one significant effect (per-protein FDR < 0.10). Representative proteins are labeled for each group, (b) Global volcano plot of all gene-protein pairs. Blue = downregulated, red = upregulated (per-protein FDR < 0.10). Black rings = self-targeted positive controls. 4,297 significant pairs, 195 self-targeted. (c) Per-protein volcano plots for four proteins with distinct regulatory profiles: CD278 (ICOS, 65 hits), CD223 (LAG3, 361 hits), CD5 (102 hits), and CD134 (OX40/TNFRSF4, 85 hits). (d) Barplot of significant regulators per surface protein (up vs. down, log scale, per-protein FDR < 0.10). (e) Gene-gene correlation heatmap for 299 genes with ≥3 significant protein effects, hierarchically clustered (Ward.D2) into 10 groups (C1-C10). (f) Mean perturbation Z-score matrix: regulator clusters (C1-C10) × protein groups. Each cluster is annotated with its dominant pathway enrichment.

Across approximately 3.5 million gene-protein pairs, 4,297 reached significance (per-protein FDR < 0.10; **Fig. 2b**). Self-targeted positive controls were enriched for positive effects, as expected for a gain-of-function screen. Proteins differed in both the number of regulators and the balance between activating and repressing regulators (**Fig. 2c, d**). CD223 (LAG-3) had the most regulators of the four shown (195 activating, 166 repressing), while CD278 (ICOS) had far fewer at a roughly even split (38 activating, 27 repressing). CD5 (39 activating, 63 repressing) and CD134 (OX40; 29 activating, 56 repressing) skewed toward repression. Because activating a gene can lower a surface protein as well as raise it, the screen measures both directions of regulation rather than induction alone.

We next grouped genes by their perturbation profiles. In contrast to the protein groups above, which cluster proteins by baseline co-expression, this analysis clusters genes by the similarity of their effects across proteins, on a subset of 299 genes with significant effects on three or more proteins. We restricted to these genes because a gene with one or two significant effects gives too sparse a profile for correlation-based clustering. We computed pairwise Pearson correlations among the 299 genes across the 201 proteins and clustered them into ten regulator clusters (C1 to C10; **Fig. 2e**). Each cluster was annotated by pathway overrepresentation analysis (MSigDB Hallmark^28^, KEGG^29^, Reactome^30^, GO Biological Process^31^, and CORUM^32^; **Methods**).

To test whether the pathway annotations matched the perturbation effects, we computed each cluster’s mean perturbation Z-score across the five protein groups (**Fig. 2f**). Because genes were clustered on their perturbation effects alone, with no pathway information, a cluster’s annotation and its effect profile come from independent sources. For most clusters the two were concordant: the pathway enrichment aligned with both the protein group most affected by the cluster and the direction of the effect. Asterisked cells in Fig. 2f mark cluster-by-protein-group combinations significant by a permutation test on the cluster labels (**Methods**), and we describe the strongest below. C7 (TNF receptor signaling) contained TNFRSF1A, TNFRSF1B, and CD27, and most strongly upregulated Activation markers (mean Z = 1.16). C8 (TCR signaling, including PIK3CA and CD81) had the strongest positive effect on Core markers (mean Z = 1.97). Because pathway annotations were assigned only after effect-based clustering, the grouping of TCR-signaling genes and their effects on Core markers were data-driven rather than annotation-driven. C3 produced the strongest repression of Core and Activation markers (mean Z = −1.95 and −1.40). This cluster contained CALM3 and NFATC2, both of which promote NFAT signaling^33,34^, and activation of either gene repressed Core and Activation markers. C9 (27 genes, protein localization to cell periphery) primarily affected Adhesion markers (mean Z = 0.49), and C4 (49 genes, calcium ion transport) produced moderate repression across several protein groups. Two protein groups showed near-zero mean Z across all clusters, for different reasons. Innate-like proteins have low baseline expression (median 1.7 versus 13.4 ADT UMI for Core), which limits the effects detectable on them. Proteins in the Other group are expressed at levels comparable to Adhesion (median 2.8 versus 3.0 ADT UMI) but each has few genes that significantly regulate it (median 1 versus 3 for Adhesion, per-protein FDR < 0.10), so their mean response stays near zero.

### Semi-NMF decomposition identifies modules of the CD4 T cell surface proteome

While individual gene-protein pairs and separate hard-clustering workflows (**Fig. 2**) reveal isolated regulatory relationships, they cannot capture overlapping gene functions or identify proteins that co-vary across multiple distinct perturbations. Matrix factorization overcomes both limitations by jointly embedding genes and proteins into a continuous space, summarizing the 201 protein columns into a small number of shared response patterns while giving each gene a continuous coefficient on each. To implement this, we factorized the gene-by-protein effect matrix (2,848 significant genes by 201 surface proteins) with semi-nonnegative matrix factorization^35^ (semi-NMF). Semi-NMF decomposes the matrix into a signed gene coefficient matrix W and a non-negative protein signature matrix H (**Fig. 3a**). The H row weights the 201 proteins and is the module’s protein signature. The W column gives every gene a signed coefficient for how strongly its perturbation activates or represses the module. Because W and H are continuous, every gene has a coefficient on all five modules and can be ranked within each, which a hard cluster label cannot represent. Rank selection by cophenetic correlation identified K = 5 modules (**Supplementary Fig. 4, Methods**).

**Figure 3:**
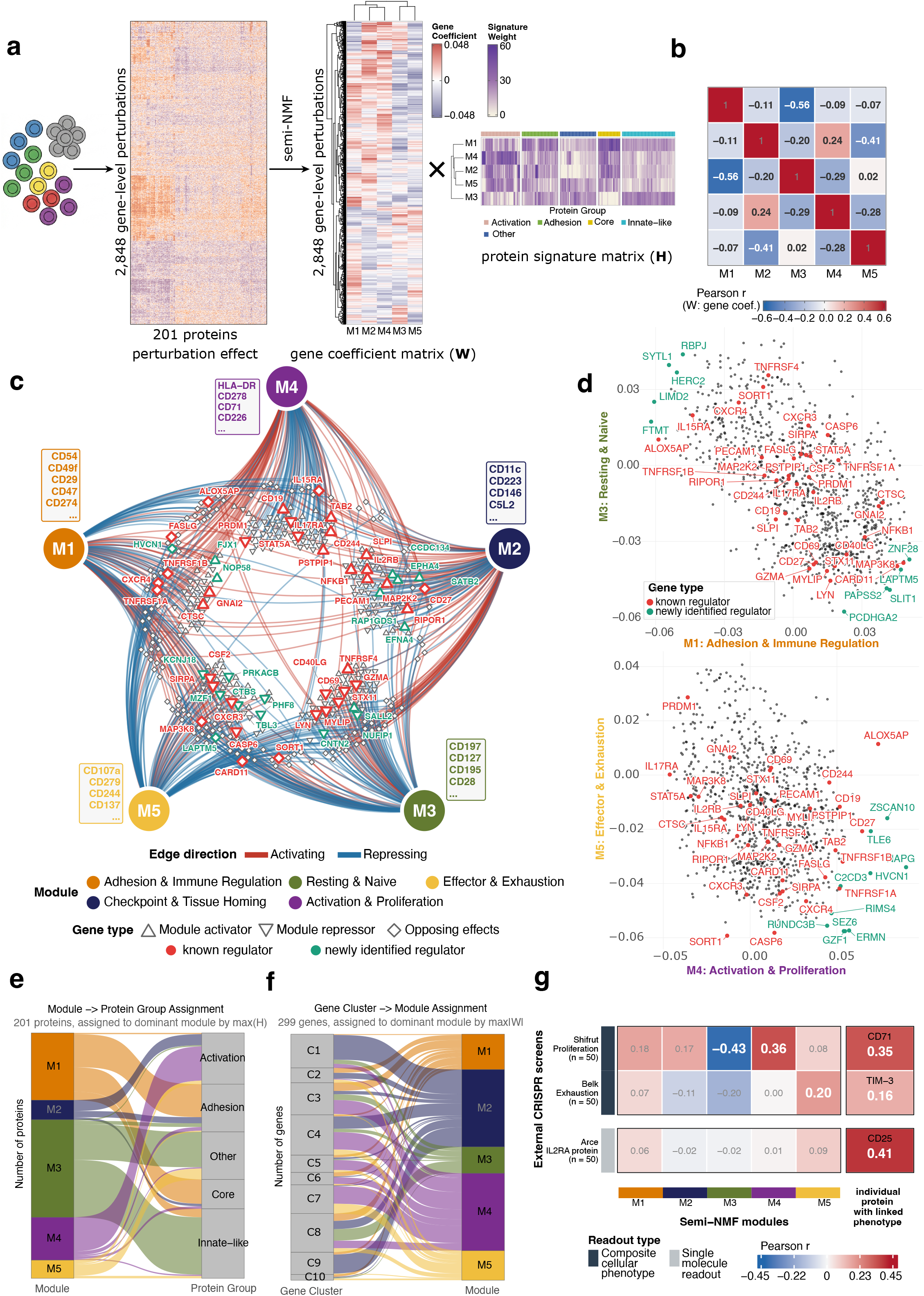
Semi-NMF Decomposition Identifies Composite Modules of the CD4 T Cell Surface Proteome. (a) Semi-NMF decomposition of the gene-by-protein effect matrix (2,848 significant genes × 201 proteins) into a signed gene coefficient matrix W (2,848 × 5) and a non-negative protein signature matrix H (5 × 201). The protein signature heatmap shows how each module loads onto the protein groups of Fig. 2a. Modules M1 to M5 are Adhesion & Immune Regulation, Checkpoint & Tissue Homing, Resting & Naive, Activation & Proliferation, and Effector & Exhaustion. (b) Pearson correlation matrix of module gene coefficients (W). (c) Network of gene perturbation effects on five T cell phenotypic modules identified by semi-NMF decomposition (k=5). Modules are positioned as a pentagon with anti-correlated pairs placed maximally apart (M1 vs M3, M2 vs M5). Each node represents a gene perturbation with large association with at least one module. Node shape encodes effect directionality (△ activator, ▽ repressor, ◇ opposing effects across modules). Edge color indicates activating (red) or repressing (blue) relationships. For activators and repressors, radial distance from a module reflects effect strength, with stronger effects positioned closer. Opposing-effect genes are positioned between their activated and repressed modules. Known regulators (bright red) and newly identified regulators (teal) are labeled. Boxed protein lists show top-loading surface markers per module from the H matrix. (d) Gene coefficients (W) for two module pairs. Top, M3 against M1; bottom, M5 against M4. Each point is a gene, placed by its coefficients on the two modules. (e) Correspondence between modules and the baseline protein groups of Fig. 2a. Alluvial plot linking each protein’s module assignment (largest H value) to its baseline co-expression group. Module assignment is associated with baseline group (permutation Z = 65.0, P < 1 × 10⁻⁴; adjusted Rand index 0.37). Per-group enrichments (Fisher’s exact): Activation to M4, 28 of 42, OR = 41.8, FDR = 3.5 × 10⁻¹⁶; Core to M1, 20 of 24, OR = 19.9, FDR = 6.6 × 10⁻⁹; Adhesion to M1, 27 of 39, OR = 10.6, FDR = 5.2 × 10⁻⁹; Innate-like to M3, 47 of 57, OR = 15.5, FDR = 7.9 × 10⁻¹⁴; Other to M3, 29 of 39, OR = 6.2, FDR = 6.5 × 10⁻⁶. (f) Correspondence between the regulator clusters of Fig. 2e and the modules. Alluvial plot linking the regulator cluster of each of the 299 genes with significant effects on three or more proteins to its dominant module (largest |W| value). Cluster membership is associated with module assignment (permutation Z = 72.0, P < 1 × 10⁻⁴; 9 of 10 clusters individually non-random, BH-adjusted P < 0.05). (g) Correlation between module gene coefficients (W) and effect sizes from three external CRISPR screens. For each screen, the top 50 SCITO-Perturb-seq significant genes ranked by external significance were used (n = 50 per screen). Right, correlation for one pre-specified single protein per screen with established relevance to the screen phenotype: CD71 for Shifrut proliferation, TIM-3 (CD366) for Belk persistence, and CD25 (IL-2Rα) for Arce IL2RA.

The five modules correspond to distinct CD4 T cell surface phenotypes, annotated from representative high-loading proteins in H (**Fig. 3a**). M1 (Adhesion & Immune Regulation) is led by the adhesion molecules CD54 (ICAM-1), CD49f (integrin α6), and CD29 (integrin β1) and the immune regulatory receptors CD274 (PD-L1) and CD47. M2 (Checkpoint & Tissue Homing) is led by CD11c, the checkpoint receptor CD223 (LAG-3), and the tissue-homing molecule CD146 (MCAM). M3 (Resting & Naive) is led by the naive and chemokine receptors CD197 (CCR7), CD127 (IL-7Rα), CD195 (CCR5), and CD28. M4 (Activation & Proliferation) is led by the activation and metabolic markers CD71 (transferrin receptor), HLA-DR, CD278 (ICOS), and CD226 (DNAM-1). M5 (Effector & Exhaustion) has a low-amplitude signature spread across effector and checkpoint markers including CD107a (LAMP-1), CD279 (PD-1), CD244 (2B4)^36^, and CD137 (4-1BB), with no single dominant protein. The maximum H value in M5 is 1.2, compared with 8.3 in M1 and 5.2 in M4, and under gene resampling M5 has the lowest signature reproducibility of the five modules (median matched H-correlation 0.63, against 0.85 to 0.93 for M1 to M4; **Supplementary Fig. 5**). We therefore do not interpret individual protein ranks within M5. We next examined the relationship among the modules by comparing their W coefficients (**Fig. 3b**). M1 and M3 are the most anti-correlated modules (r = −0.56) and M2 and M5 the next (r = −0.41), so genes that activate one member of either pair tend to repress the other. The remaining module pairs are weakly correlated (|r| ≤ 0.30).

To translate these module relationships into a global gene-level regulatory map, we mapped gene perturbations to modules by building a network that links each gene to the modules it strongly activates or represses (**Fig. 3c, Methods**). Many known regulators connect to a single module in the expected direction. CD69, CD40LG, and GZMA all repress M3 (Resting & Naive). CD69 is the earliest surface marker of T cell activation and retains activated T cells in lymphoid organs by opposing S1PR1^37^. GNAI2 activates M1 (Adhesion & Immune Regulation) and encodes the Gαi2 subunit required for chemokine receptor signaling and T cell homing to lymph nodes^38^. EPHA4 and its ligand EFNA4 each activate M2 (Checkpoint & Tissue Homing). EPHA4 promotes CD4 T cell chemotaxis through activation of VAV1, LCK, and FYN^39^, and finding a receptor and its ligand on the same module is an internal consistency check on the decomposition. PRDM1 (BLIMP-1) represses M4 (Activation & Proliferation), consistent with its direct transcriptional repression of IL-2 and FOS in CD4+ T cells that limits clonal expansion following TCR stimulation^40^. Note that these associations were not revealed in examining each pair of sgRNA and surface protein expression individually.

Because W is continuous and signed, a gene can load positively on one module and negatively on another (**Fig. 3d**). At the 90th percentile of |W| (**Methods**), 294 of the 2,848 genes (10.3%) have a strong coefficient on two or more modules, and 266 (9.3%) have a positive coefficient on one module and a negative coefficient on another. Several of these multi-module regulators have established mechanisms. CARD11 has a positive M1 coefficient and a negative M3 coefficient (**Fig. 3d, top**), and it scaffolds the CBM signalosome that drives NF-κB activation downstream of the TCR^41^. LAPTM5 also has a positive M1 and a negative M3 coefficient and lowers surface TCR by routing CD3ζ for lysosomal degradation^42^. PRDM1 has a negative M4 and a positive M5 coefficient (**Fig. 3d, bottom**). TNFRSF1A and TNFRSF1B have positive M4 and negative M5 coefficients. These M5 coefficient signs are reproducible under gene resampling, unlike the M5 protein ranks (positive for PRDM1 in 87% of replicates, negative for TNFRSF1A and TNFRSF1B in 87% and 81%).

Semi-NMF jointly models gene and protein structure in the perturbation effect matrix, so we compared the modules with the baseline protein groups (**Fig. 2a**) and the regulator clusters (**Fig. 2e**) to ask whether the decomposition recovers those groupings or adds to them. In a held-out reconstruction benchmark, the modules predicted masked entries of the effect matrix more accurately than either the protein groups or the regulator clusters (**Methods**), so the continuous factorization captures effect-matrix structure that the hard groupings miss. To compare the partitions directly, we assigned each protein to the module with its largest H value and compared this partition with the five protein groups defined by baseline co-expression (**Fig. 3e**). The two partitions overlap but are not identical (adjusted Rand index 0.37). Where the partitions agree, the modules recover baseline structure, as for the Activation proteins mapping to M4 (28 of 42, FDR = 3.5 × 10⁻ ¹⁶). Where they diverge, the modules group proteins that baseline co-expression separates. M1 combines the Core and Adhesion groups (20 of 24 Core and 27 of 39 Adhesion proteins map to M1; FDR = 6.6 × 10⁻ ⁹ and 5.2 × 10⁻ ⁹). The Innate-like and Other groups both map to M3 (47 of 57 and 29 of 39; FDR = 7.9 × 10⁻ ¹⁴ and 6.5 × 10⁻ ⁶). These proteins are expressed at low levels in resting cells and change little under perturbation (**Fig. 2f**), so their assignment to M3 follows from a weak response rather than a shared program, while M3’s signature is set by its naive and resting markers such as CD197 (CCR7) and CD127. M2 is the strongest case of the modules grouping proteins that baseline co-expression does not. M2’s high-loading proteins are checkpoint and tissue-homing markers such as CD223 (LAG-3), CD11c, and CD146, which the baseline clustering places in different groups (**Supplementary Table**). Baseline co-expression therefore does not assemble this set, and it appears only in the perturbation response. M2 is also reproducible under gene resampling (median matched H-correlation 0.91; **Supplementary Fig. 5**).

The gene coefficients W relate the modules to the regulator clusters of Fig. 2e (**Fig. 3f**). Where a cluster’s genes act on one phenotype, it maps to a single module, with C9 (protein localization to cell periphery) assigning 85.2% of its genes to M2, and C1 (ABC transporters) assigning 73.2% to M2. Where a cluster’s genes act on multiple phenotypes, their module coefficients were distributed across multiple modules. C7 (TNF receptor signaling) splits between M4 (60.0%) and M5 (34.3%), consistent with TNF superfamily costimulation engaging both proliferation and terminal differentiation. C8 (TCR signaling) has no majority module and spreads across M1 to M4, consistent with the broad effects of TCR engagement. The regulator clusters assign each gene to one group. The module coefficients additionally show that some of those groups, such as C7 and C8, act across several phenotypes rather than one.

Having established that the modules capture structure within the SCITO-Perturb-seq dataset, we next tested whether they correspond to independently measured T cell functions. We therefore compared the module gene coefficients with effect sizes from three published T cell CRISPR screens (**Fig. 3g**). Shifrut et al.^9^ screened for regulators of proliferation by CFSE dilution after TCR stimulation, Belk et al.^11^ for regulators of persistence under chronic antigen stimulation, and Arce et al.^14^ for regulators of IL2RA surface expression by FACS-based sorting after knockout. For each screen we ranked the 2,848 genes with at least one significant protein effect by their significance in that external screen, and took the top 50. For the Shifrut proliferation screen, M4 (Activation & Proliferation) was positively correlated (r = 0.36, p = 0.010) and M3 (Resting & Naive) was anti-correlated (r = −0.43, p = 0.002), so genes that drive proliferation repress the resting surface phenotype. M3’s anti-correlation was stronger than that of CCR7, the canonical naive marker (r = −0.30, p = 0.034), whereas M4 tracked proliferation about as well as CD71, the single marker most closely linked to this screen (r = 0.35, p = 0.012). The resting module therefore captures the proliferation axis better than any single marker, a multi-protein relationship that no individual protein reproduces. For the Belk persistence screen, M5 (Effector & Exhaustion) had the highest module correlation (r = 0.20, p = 0.16) and exceeded the exhaustion marker TIM-3 (r = 0.16, p = 0.27), although neither correlation reached significance at this sample size. For the Arce IL2RA screen, CD25 was strongly correlated (r = 0.41, p = 0.003) while all five modules remained near zero (|r| ≤ 0.10, all p > 0.5). M3’s anti-correlation with proliferation is the strongest cross-screen signal and is a directional relationship that no single marker reproduces, while the single-protein IL2RA screen tracks CD25 and not the modules. The modules therefore correspond to multi-protein phenotypes rather than to single gene-protein associations.

## DISCUSSION

Pooled CRISPR screens in primary T cells have been restricted to either profiling the global transcriptome or isolating cells by a single functional phenotype-such as an individual surface protein-neither of which directly measures the coordinated expression of surface markers that define T cell identity. SCITO-Perturb-seq closes this gap by combining pooled CRISPR activation with combinatorial-indexed single-cell cytometry. This framework enables genome-wide CRISPR activation to be directly mapped to high-dimensional surface proteomes at a fraction of the per-cell cost of transcriptome sequencing. In 3.6 million primary human CD4 T cells across 18,325 gene activations, 16% of perturbed genes alter at least one of 201 surface proteins. Semi-NMF decomposition of the perturbation effect matrix identifies five modules of the CD4 T cell surface proteome, each pairing a set of gene perturbations with a surface-protein signature that maps to a known T cell phenotype. These modules group proteins by their shared response to perturbation, including proteins not co-expressed in unperturbed cells.

While platforms like ECCITE-seq^22^ and Perturb-CITE-seq^24^ couple single-cell RNA and surface protein measurements with CRISPR perturbations, they remain restricted to small, targeted gene panels. SCITO-Perturb-seq achieves genome-wide perturbation scale while simultaneously profiling hundreds of surface proteins across millions of individual cells. This leap in throughput is driven by combinatorial indexing, which expands cell capacity per microfluidic channel roughly 20-fold, while a defined antibody panel minimizes sequencing overhead. Direct protein measurement is biologically essential, as it captures surface phenotype changes driven by both transcriptional and post-translational regulation. Furthermore, a CRISPRa modality is well suited to profiling protein expression at a single, static timepoint. Because many surface proteins are highly stable and turn over slowly, a CRISPR knockout or CRISPRi screen requires waiting days or weeks for pre-existing proteins to naturally degrade, which may reduce the power of the study at an early harvest window. Conversely, CRISPRa actively drives the immediate synthesis and trafficking of new protein and thus bypasses the lag of baseline protein stability, providing a highly sensitive readout of genetic regulation at a single snapshot in time.

While SCITO-Perturb-seq provides an unprecedented map of T cell regulation, several structural parameters define the scope of this initial dataset. First, our map utilizes cells from a single donor to establish a robust baseline, meaning donor-to-donor variation in these regulatory networks remains a subject for future cohort-scale studies. Second, because CRISPRa induces robust gene overexpression, the phenotypes documented here represent the cellular response to elevated expression rather than baseline endogenous regulation. Third, our measurements capture a highly defined single snapshot following T cell activation. While this strategically captures a broad swath of the proteome, it does not resolve the distinct temporal dynamics that characterize early versus late-stage activation markers. Finally, our phenotypic lens is focused on a curated 201-protein panel optimized for T cell lineage, activation, and exhaustion markers. Within this panel, a small subset of chemokine receptors, such as CCR7, may exhibit conservative effect sizes due to the standard 4°C staining workflow required for large-scale single-cell processing.

Looking forward, the core architecture of SCITO-Perturb-seq provides a highly scalable blueprint designed to adapt alongside the rapidly evolving landscape of single-cell genomics. While our initial 10-pool implementation was bounded by the microfluidic architectures available at the time, the platform is perfectly positioned to leverage next-generation single-cell chemistries. By integrating with newer fixed-cell or multi-channel loading formats, this strategy seamlessly scales to 96-well configurations—unlocking simultaneous transcriptomic profiling, high-throughput multi-donor multiplexing, and dense kinetic time-courses within a single experiment. Beyond CD4+ T cells, this combinatorial-indexing architecture serves as a universal plug-and-play blueprint across primary immune populations, easily accommodating CRISPRi or base editing to chart necessity and precise allelic effects. Ultimately, by mapping 201 surface proteins against 18,325 gene activations, this work delivers the first comprehensive, protein-level atlas of human genetic perturbations. This genome-wide blueprint provides an enduring foundation for understanding immune cell state and function, serving as a powerful discovery engine to accelerate mechanistic immunology and therapeutic target identification.

## Methods

### Cell culture and lentiviral production

CD4+ T cells were isolated using EasySep magnetic selection following the manufacturers’ recommended protocol (STEMCELL Technologies, catalog no. 17951). CD4+ T cells were cultured in X-VIVO 15 (Lonza Bioscience, catalog no. 04-418Q) supplemented with 5% fetal bovine serum (FBS), 55 mM 2-mercaptoethanol, 4 mM N-acetyl L-cysteine, and 500 IU/ml of recombinant human IL-2 (Amerisource Bergen, catalog no. 10101641). Primary T cells were activated using anti-human CD3/CD28 CTS Dynabeads (Fisher Scientific, catalog no. 40203D) at a 1:1 cell:bead ratio at 10^6^ cells/ml.

Lenti-X HEK293T cells (Takara Bio, catalog no. 632180) were maintained in high-glucose Dulbecco’s modified Eagle’s medium with GlutaMAX (Fisher Scientific, catalog no. 10566024), supplemented with 10% FBS, 100 U/ml of penicillin/streptomycin (PenStrep; Fisher Scientific, catalog no. 15140122). Lentivirus was produced by co-transfecting HEK293T cells with transfer plasmids and standard packaging vectors using Lipofectamine 3000 transfection reagent (Fisher Scientific, catalog no. L3000075) as described previously^10^.

### Genome-wide SCITO-Perturb-seq library design and cloning

The guide sequences (two guide sequences per gene) were selected from the genome-wide CRISPRa (Calabrese A, catalog no. 92379 and Calabrese B, catalog no. 92380) libraries considering the expression preference in primary human T cells in the previous paper^10^. To achieve combinatorial indexing of guides, all guides are distributed roughly equally to ten different bins with no overlaps among pools. Non-targeting control guides were randomly sampled from two different Calabrese sets (A and B), which accounted for 5% of the total library size in each bin. Importantly, to minimize the loss of signal of particular gene targets, we distributed two guide sequences (targeting the same gene) into two different bins.

Three plasmids (dCas9 vector, sgRNA cloning vector and activation effector vector) were used in this study. For dCas9-VP64 expression we used Lenti-SFFV-mCherry-2A-dCas9-VP64 (pZR112, Addgene #180263) as described previously^10^. The Lenti-PCP-P65-HSF1 vector (pZR232) was constructed by replacing mCherry-dCas9-VP64 in pZR112 with PCP-P65-HSF1-2A-PuroR from pXPR_502 (Addgene #96923). The sgRNA lentiviral expression vector (pZR235) was derived from the parental CROPseq-Opti (Addgene #106280). First puromycin resistance was replaced with EGFP. Second the sgRNA scaffold was replaced with a modified version of the pXPR_502 scaffold, only containing the PP7 stem loops (and not MS2). Oligos encoding the sgRNA library were synthesized as an oligonucleotide pool (IDT) with flanking sequences compatible with Golden Gate Cloning (BsmBI-v2, New England Biolabs, catalog no. E1602L). Oligo pools were amplified, digested with BsmBI, and ligated into pZR235 to make genome-wide guide libraries.

### SCITO-Perturb-seq screening and droplet GEM generation

The selection of time points for co-transducing the lentiviral vector was the main goal of our experiment. After multiple rounds of optimization, we were able to obtain optimal time points and good viral titers to achieve target MOI of ∼0.3 of sgRNA vector. Primary human CD4+ T cells were isolated from one healthy PBMC donor. One day after activation with CD3/28 beads, T cells from a human blood donor were infected with 4% v/v concentrated dCas9-VP64 lentivirus (pZR112) and the 2% v/v guide vector (pZR235). Two days after activation, T cells were infected with 2% v/v of the activation helper vector. Cells were maintained at a viability of >90%, and a density of ∼ 1 × 10^6^ cells/ml for the course of the experiment. Three days after T cell activation, a fresh medium with IL-2 (final concentration 500 IU/ml) and puromycin (final concentration 1.5 μg/ml) was added to bring cells to 3 × 10^5^ cells/ml. Cells were split two days later and fresh medium with IL-2 was added to bring cells to 3 × 10^5^ cells/ml. After two days, fresh medium without IL-2 was added to bring the concentration to 10^6^/ml. Eight days after initial activation, cells were harvested, centrifuged at 500g for 5 min, and resuspended at 2 × 10^6^ cells/ml X-VIVO 15 without supplements. The following day, cells were restimulated. The next day, cells were sorted to near purity by FACS (FACSAria2, BD Biosciences), using GFP as a marker for sgRNA vector transduction and mCherry for the dCas9 expression.

Counted cells (Countess II, ThermoFisher) were prepared by resuspension in 1X PBS with 0.04% BSA as detailed in the 10x Genomics Single Cell Protocols Cell Preparation Guide (10x Genomics, CG00053 Rev C). 5 × 10^5^ cells were then loaded into each channel using the Chromium X Controller (10x Genomics) with Chromium Single-Cell 3′ HT Gel Beads v3.1 (10x Genomics, PN-1000348 and PN-1000349) across 22 “lanes”/GEM groups” following the 10x Genomics user guide (CG000417).

### SCITO-Perturb-seq library preparation and sequencing

For preparation of ADT expression and sgRNA libraries, samples were processed according to 10x Genomics Chromium Next GEM Single Cell 3ʹ HT Reagent Kits v3.1 (Dual Index) with Feature Barcode technology for Cell Surface Protein (CG000417). After cDNA amplification step, final ADT library was amplified using one step PCR with Dual Index Kit NT Set A. For sgRNA library, stepwise sgRNA enrichment was performed as previously described^43^. For sequencing, ADT and sgRNA libraries were pooled at a 2:1 ratio to maintain the library complexity. Libraries were sequenced on a NovaSeq 6000 (Illumina) platform.

### Quantification and statistical analysis

#### Alignment, droplet deconvolution and guide assignment

Cell Ranger 6.0.0 software (10x Genomics) was used for alignment of ADT reads to the custom antibody barcode sequences. Reads from the sgRNA libraries were mapped with custom sgRNA sequences using modified Drop-seq alignment pipeline (https://github.com/broadinstitute/Drop-seq/blob/master/doc/Drop-seq_Alignment_Cookbook.pdf). Raw guide UMI counts were merged from all GEMs and droplet-level multiplets (multiple cells in one droplet but can be deconvoluted using combinatorial barcodes) were deconvoluted as previously described^27^. The guide assignment was performed using a simple procedure with normalized counts of sgRNAs. For each SCITO-Perturb-seq pool, treated as a batch, the sgRNA count matrix was rescaled between 0 and 1 by dividing each sgRNA count by its maximum value. Guide assignments were determined by selecting the sgRNA with the highest count for the cells in each batch.

### Quality control and ADT count normalization

Cells were filtered to keep those with total ADT UMI counts between 300 and 15,000, and no single protein marker exceeding 5,000 counts. Batch 8 was excluded from analysis due to quality concerns, leaving 9 batches (Batches 1-7, 9-10). This resulted in 201 surface protein measurements and 7 isotype controls across 3,134,864 cells passing filters.

Normalization was performed independently within each batch using a Poisson generalized linear model (GLM) framework to regress out technical covariates. For each batch, isotype control counts (n = 7) were log-transformed (log (count + 1)) and reduced to three principal components via PCA. For each protein marker, a Poisson GLM was fit using the first two isotype PCs and log-transformed library size (log (total ADT counts + 1)) as covariates. Deviance residuals from the fitted model were extracted as the normalized expression values, which account for both ambient antibody background (captured by isotype PCs) and cell-level capture efficiency (captured by library size). Deviance residuals were then z-scored (zero mean, unit variance) within each batch and concatenated across all 9 batches to produce the final normalized dataset.

### Perturbation effect estimation

For the primary gene-level analysis, all cells carrying any sgRNA targeting the same gene were pooled. Genes with fewer than 20 assigned cells were excluded (419 genes removed, 2.4%), resulting in 17,336 genes with a median of 150 cells per gene (range: 20-1,266). Non-targeting control (NTC) cells (n = 26,602) served as the reference population. A secondary sgRNA-level analysis was performed in parallel, in which each of the 28,104 individual sgRNAs (median 88 cells; range: 1-1,237) was analyzed independently for use in concordance validation.

For each gene-protein pair, the normalized expression difference was computed as the difference between the mean deviance-residual expression in gene-perturbed cells and the mean in NTC cells. Standard errors were calculated using the Welch approximation: SE = √ (s²_gene/n_gene + s²_NTC/n_NTC), where s² denotes sample variance with Bessel’s correction. Z-scores were computed as normalized expression difference / SE, and two-sided p-values were obtained from Welch’s t-test using the Satterthwaite approximation for degrees of freedom. Effect sizes at the sgRNA level were computed identically, substituting individual sgRNA cell populations for gene-level pools. p-values from the gene-level Welch’s t-test were adjusted using the Benjamini-Hochberg procedure applied independently within each protein (across 17,336 genes per protein).

### Self-targeted validation

A curated mapping between the 201 surface protein markers (excluding 7 isotype controls) and their encoding gene(s) was constructed manually from established nomenclature (e.g., CD25 → IL2RA, CD38 → CD38, CD62L → SELL). For proteins recognized by antibodies targeting products of multiple genes (e.g., HLA-ABC → HLA-A, HLA-B, HLA-C; CD3 → CD3D, CD3E, CD3G), all corresponding genes were included, resulting in 234 total gene-to-marker mappings across 177 unique proteins.

To assess assay sensitivity, we identified self-targeted gene-protein pairs in which the perturbed gene encodes the measured surface protein (e.g., CRISPRa of CD38 measured against CD38 protein), yielding 195 pairs (185 unique genes, 177 unique proteins) after cell-count filtering. Because the gene-to-protein map is many-to-many, we reduced these to one entry per gene and one entry per protein, selecting the pair with the largest |z-score| in each pass, giving a deduplicated set of 172 pairs. This 172-pair set was used for the self-targeted summary statistics and is shown in both the heatmap (Fig. 1c) and the volcano plot (Fig. 1d). Because CRISPRa induces transcriptional activation, the expected ground truth for self-targeted pairs is positive normalized expression difference. Statistical significance was assessed using per-protein FDR (BH correction across all 17,336 genes per protein).

For self-targeted genes with exactly two independent sgRNAs present we compared the normalized expression difference of each sgRNA against the cognate protein. This resulted in 122 self-targeted gene-protein pairs from 112 unique genes. Concordance was defined as both sgRNAs producing a positive effect on the cognate protein. Spearman correlation (ρ) was computed between paired sgRNA normalized expression difference values.

### Split-half reliability

To evaluate the reproducibility of normalized expression difference estimates, we performed split-half reliability analysis. For each gene, cells were randomly partitioned into two independent halves (A and B). Half B served as held-out validation (using 100% of its cells), while half A was progressively downsampled at fractions of 10%, 25%, 50%, 75%, and 100%. Gene-level normalized expression difference was computed independently for each half, and Pearson correlation between the two halves was calculated across significant pairs (per-protein FDR < 0.10), and self-targeted pairs. This procedure was repeated across 10 independent random splits, and results were reported as mean ± standard deviation. Genes with fewer than 10 cells in either half after splitting were excluded. The correlation at 100% of half A represents the reliability ceiling, which is the maximum achievable correlation between independent cell samples, bounded by measurement noise rather than sample size.

### Cross-study validation

To validate SCITO-Perturb-seq CD25 (IL2RA) effect estimates against published screens, we compared our gene-level CD25 Z-scores against two loss-of-function datasets: (1) IL2RA trans-effect Z-scores from a CRISPRi perturb-seq screen (Zhu et al., resting condition); and (2) IL2RA log fold change (LFC) values from a CRISPR knockout FACS screen in stimulated T effector cells (Arce et al., Stim72hr condition). For the Arce dataset, genes were filtered to those reaching FACS significance (FDR < 0.05). Pearson correlations were computed on the three-way gene intersection across all three datasets. In the figure, Zhu Z-scores are negated so that positive values indicate positive regulators, matching the convention of the original Arce LFC values and SCITO-Perturb-seq Z-scores, and positive slopes therefore indicate agreement across screens.

### Protein-protein co-expression and perturbation Z-score heatmap

To characterize baseline surface protein co-expression, we extracted all non-targeting control (NTC) cells (n = 26,602) from the deviance-normalized expression matrix. Pearson correlation coefficients were computed across all 201 surface proteins on the NTC expression profiles, yielding a 201 × 201 protein-protein correlation matrix. Proteins were hierarchically clustered using Ward.D2 linkage on a distance metric of 1 − r, and the resulting dendrogram was cut at k = 5 to define protein groups.

For the protein × gene (P×G) heatmap (Fig. 2a, right), we selected the 2,848 genes with at least one significant protein effect at per-protein FDR < 0.10. Z-scores (LFC/SE) rather than raw log fold changes were used for visualization, as Z-scores are confidence-weighted and downweight noisy estimates from genes with few cells. Genes were hierarchically clustered (Ward.D2 on Euclidean distance of the Z-score matrix).

### Gene-gene correlation and clustering

To group genes by the similarity of their effects across proteins, we selected 299 genes with ≥3 significant protein effects (per-protein FDR < 0.10). Self-targeted gene-protein pairs were masked (set to NA, then zero-filled for clustering) to prevent on-target effects from dominating the correlation structure. Pearson correlation was computed on the masked Z-score profiles across 201 proteins, producing a 299 × 299 gene-gene correlation matrix. Genes were hierarchically clustered using Ward.D2 linkage on the correlation-based distance (1 − r) and partitioned into k = 10 clusters. The optimal k was evaluated by silhouette analysis over k = 5-20, with k = 10 selected to balance resolution and interpretability (Fig. 2e).

Each regulator cluster was characterized by its mean perturbation Z-score across the five NTC-derived protein groups (Fig. 2f). Intra-cluster correlation was computed as the mean pairwise Pearson correlation among cluster members. Pathway enrichment was performed using overrepresentation analysis (clusterProfiler^44^ in R) against multiple databases: MSigDB Hallmark gene sets, KEGG (via MSigDB), Reactome, Gene Ontology Biological Process (GO:BP), and CORUM protein complexes.

Gene symbols were mapped to Entrez IDs using org.Hs.eg.db, and the background universe included all 17,336 tested genes with valid Entrez mappings. Gene sets were filtered to 5-200 members, and a minimum overlap of 2 genes was required. Enrichment significance was assessed at BH-adjusted p < 0.25. Where automated annotation was suboptimal, manual corrections were applied based on inspection of the top enrichment terms and the constituent gene perturbation profiles.

### Mean perturbation Z-score for each regulator cluster (C1 to C10) and protein group

Z-scores come from a Welch t-test of perturbed against NTC cells and are precision-weighted, so genes tested in more cells contribute more to the mean. The Z-score is not itself an effect size. Cells marked with an asterisk are significant at permutation BH-FDR < 0.05, with gene-to-cluster labels shuffled 10,000 times. The same strong-versus-weak pattern holds when each cell is recomputed from the raw normalized expression difference in place of the Z-score, so the cluster-group associations are not an artifact of precision weighting.

### Module identification via semi-NMF

To decompose surface protein changes across perturbations, we applied semi-nonnegative matrix factorization (semi-NMF) to the gene-level perturbation effect size matrix. Unlike standard NMF, which requires non-negative input data, semi-NMF allows negative values in the input matrix and in the gene coefficient matrix W. This formulation is well-suited for perturbation data where gene activation can either increase or decrease surface protein expression relative to non-targeting controls.

Starting from the gene-level z-score effect matrix (17,336 genes × 201 surface proteins), we first filtered to genes with at least one significant protein association (per-protein FDR < 0.1 by Benjamini-Hochberg correction), yielding 2,848 genes. To prevent cis-regulatory effects from dominating module structure, we masked self-targeted gene-protein pairs by setting their values to zero prior to factorization (195 pairs masked). Remaining missing or non-finite values were set to zero. We evaluated four input matrix variants in a systematic comparison: perturbation effect sizes z-score versus raw effect sizes, each with or without cis-pair masking. All four variants were subjected to identical rank scans (K = 5-25, 10 runs per rank, 100 maximum iterations). Factorization stability, as measured by cophenetic correlation, was comparable across variants, with all four identifying K = 5 as the most stable rank. We selected the z-score, cis-masked variant (value range: [−17.34, 9.82]) for downstream analysis, as z-scores provide a standardized scale that accounts for estimation uncertainty and cis-masking prevents trivially strong self-regulatory signals from driving module structure.

The semi-NMF decomposition factorizes the effect matrix X (genes × proteins) into X ≈ WH, where the gene coefficient matrix W (genes × K) quantifies each gene perturbation’s contribution to each of K phenotypic modules, and the protein signature matrix H (K × proteins) defines the characteristic surface protein expression pattern of each module. The semi-NMF formulation constrains H ≥ 0 while allowing negative entries in W, allowing gene perturbations to either activate (positive coefficients) or oppose (negative coefficients) a module’s protein signature. Non-negativity on H keeps each protein signature an additive combination of proteins. A signed W is required to represent repression, because under a non-negative constraint on W the coefficients of known repressors such as PRDM135,36, CALM3, and NFATC2 collapse toward zero. We used the semi-NMF implementation from the hNMF R package (v1.0) with random initialization.

To determine the optimal number of modules K, we performed an extended rank scan from K = 3 to K = 10 with 20 replicate runs per rank and 500 maximum iterations per run (seed = 43). We evaluated cophenetic correlation (measuring factorization stability across runs), dispersion, residual sum of squares, explained variance, and silhouette width. K = 5 was selected as the optimal rank, because cophenetic correlation was maximal at K = 5, indicating that K = 5 produced the most reproducible factorization. Explained variance increased from K = 3 through K = 10, but the marginal gain diminished beyond K = 5. We additionally compared K = 5 and K = 7 models by GO biological process and KEGG pathway enrichment analysis and by overlap with curated T cell marker gene sets (TCR signaling, T cell activation, exhaustion, cytokine signaling, costimulation, coinhibition, Treg, Th1, Th2, Th17, and migration/adhesion). Both ranks captured major T cell biology, but K = 5 showed more interpretable, non-redundant modules with stronger enrichment signals per module. The K = 5 decomposition was therefore used for all subsequent analyses.

To confirm that a signed W is required to represent repression, we fitted standard NMF (Lee and Seung multiplicative update, NMF R package) at K = 5 to the full input matrix. The matrix was shifted to non-negative before fitting by subtracting the global minimum. Under the non-negative constraint, W values for PRDM1, CALM3, and NFATC2 were all ≤ 5.5 × 10⁻⁴ across all five modules. Under semi-NMF, the same genes carry negative W coefficients on M4 (PRDM1 = −0.036, CALM3 = −0.027, NFATC2 = −0.009, L2-normalized), consistent with their roles as transcriptional repressors of T cell activation.

### semi-NMF module characterization

For visualization and cross-module comparison, we L2-normalized the columns of W (dividing each column by its Euclidean norm) and transferred the scaling factors into H by multiplying H by the diagonal matrix of column norms. This normalization places all module loadings on a common scale, enabling a principled threshold for identifying genes with strong module associations. Gene-module associations were defined using the 90th percentile of absolute L2-normalized W values as the significance threshold. Genes were classified as module activators (positive W loading above threshold), module repressors (negative W loading below negative threshold), or genes with opposing effects (exceeding threshold in both positive and negative directions across different modules).

For the network visualization, category 1 (module-defining) selected the top 50 activators and top 50 repressors per module ranked by W value in that module. Category 2 (module-specific) selected the 10 highest-magnitude genes per module among those with exactly one module above threshold. Category 3 (opposing-effect) identified genes with at least one positive and one negative association above threshold, ranked by the product of absolute positive and negative loading magnitudes, and retained the top 100. Known regulators and newly-identified regulators passing threshold were added if not already selected. For each module, surface proteins were selected from the top ten by H loading among proteins primarily assigned to that module, choosing proteins with established surface expression in the corresponding T cell state (up to five proteins per module).

Gene Ontology (GO) biological process and KEGG pathway enrichment analyses were performed for each module using clusterProfiler^44^. For each module, genes with W loadings exceeding the mean plus one standard deviation were tested against the background of all 2,848 filtered genes, using Benjamini-Hochberg correction (FDR < 0.05). Separately, negatively loaded genes (below the mean minus one standard deviation) were tested when at least 20 such genes were present in a module. Gene symbols were mapped to Entrez IDs for KEGG analysis using org.Hs.eg.db.

### Module stability assessment

Each module’s protein signature stability was assessed by a gene-resampling bootstrap with 20 replicates. In each replicate, 80% of the 2,848 filtered genes were sampled without replacement. Semi-NMF was re-fitted at K = 5 using 20 runs and 500 maximum iterations per run with random initialization (hNMF R package). The resulting H matrix was aligned to the reference H from the full-data fit by solving the linear sum assignment problem on the K × K matrix of inter-module Pearson correlations (Hungarian method, clue R package^45^). The matched H-correlation for each module is the Pearson correlation between the reference H row and the aligned bootstrap H row across all 201 proteins. Results are reported as median [IQR] across 20 replicates. Modules with median matched H-correlation below 0.70 were treated as having low protein signature reproducibility.

### Held-out reconstruction benchmark

To compare the predictive accuracy of semi-NMF against the protein groups and regulator clusters from Fig. 2, we ran a held-out reconstruction benchmark with 5 folds × 5 random seeds (25 evaluations). In each evaluation, 20% of non-zero entries in the input matrix (2,848 genes × 201 proteins) were randomly masked. Three methods were fitted on the observed entries and used to predict the masked entries. Semi-NMF K = 5 replaced masked entries with zero before fitting. Protein-group mean predicted each entry as the row-mean of that gene’s effects across all proteins in the same NTC co-expression group. Gene-cluster mean predicted each entry as the column-mean of all other genes in the same C1-C10 regulator cluster (leave-one-out), applied only to the 299 genes with a cluster assignment. Reconstruction accuracy was measured as root mean squared error on the masked entries only.

### Module concordance with protein groups and regulator clusters

To test whether module assignments are consistent with the protein groups and regulator clusters of Fig. 2, we used a permutation-based between-group sum of squares test. The test statistic is the weighted between-group sum of squares. For each group, the squared Euclidean distance between the group mean loading vector and the grand mean is computed across all module dimensions, multiplied by group size, and summed over all groups. For protein groups, the test was applied to the transposed H matrix (201 proteins × 5 modules) with the five NTC co-expression groups as labels.

For regulator clusters, the test was applied to the W matrix (299 genes × 5 modules) with the C1–C10 cluster labels. Each protein was assigned to the module with its largest H value. Each gene was assigned to the module with its largest |W| value. The null distribution was generated by permuting group labels 10,000 times with loadings fixed. Per-group and per-cluster significance used the same permutation null applied to each group separately, with Benjamini–Hochberg correction. The adjusted Rand index between the protein module partition and the NTC baseline co-expression groups was computed using the adjustedRandIndex function from the mclust R package^46^.

### Validation against external CRISPR screens

We compared module gene coefficients (columns of W) against effect sizes from three published T cell CRISPR screens: Belk et al. (knockout screen for exhaustion in vivo), Shifrut et al. (knockout screen for proliferation), and Arce et al. (knockout screen for IL2RA surface protein by FACS, stimulated T effector condition). For each external screen, the top 50 genes ranked by external significance among genes with at least one significant protein association in SCITO-Perturb-seq (per-protein FDR < 0.1) were selected, resulting in a uniform gene set size across all comparisons. Pearson correlations were computed between each module’s gene coefficient vector and the external screen effect sizes across the selected genes.

Effect sizes were adjusted per screen to account for differences in perturbation modality. No adjustment was applied to Belk et al. Positive log fold change in Belk et al. reflects knockout enrichment under chronic stimulation, indicating the gene normally drives exhaustion. Overexpression of such genes in SCITO-Perturb-seq (CRISPRa) would be expected to increase exhaustion-associated surface markers, so positive correlations reflect concordance. For Shifrut et al., we negated the MAGeCK^47^ positive-selection log fold change so that positive values correspond to genes whose overexpression would be expected to increase proliferation, matching the CRISPRa direction. For Arce et al., we used the raw MAGeCK negative-selection log fold change without sign adjustment, consistent with the loss-of-function sign convention used in Fig. 1g, where positive values indicate genes whose knockout decreases IL2RA surface expression (positive regulators of IL2RA).

Overexpression of these genes in SCITO-Perturb-seq (CRISPRa) would therefore be expected to increase CD25 expression, so positive correlations reflect concordance. As an additional comparison, we computed Pearson correlation between each external screen and the perturbation Z-score of a single surface protein with established biological relevance to the screen phenotype: TIM-3 (CD366) for exhaustion, CD71 (transferrin receptor) for proliferation, and CD25 (IL-2Rα) for IL2RA expression.

## DATA AVAILABILITY

All genome-wide pooled CRISPR screen data have been deposited at the National Center for Biotechnology Information (NCBI) Gene Expression Omnibus (accession number GSE330295).

## CODE AVAILABILITY

Custom codes to reproduce the findings of this study, in addition to analyzed Perturb-seq data, are deposited at Github (https://github.com/bjhwang113/SCITO-Perturb-seq).

## ACKNOWLEDGMENTS

We thank Z. Steinhart and R. Schmidt for the plasmid construction and all the Ye lab members and Hwang lab members for their critical comments and helpful discussions. We thank sequencing support from UCSF CAT members including Eric Chow; UCSF Flow Cytometry Core. This work was supported by the National Research Foundation of Korea (NRF) grant funded by the Korea government (MSIT) (RS-2023-00276271, RS-2025-17992968, RS-2025-02214844). C.J.Y. is supported by an NIAID R01 (R01AI171184), NIDCR U01 (U01DE028891), NIAID P01 (P01AI172523), and NHGRI UM1 (UM1HG012076). Additional support comes from NIAMS (P30AR070155), the Department of Defense, the Arc Research Institute, Parker Institute for Cancer Immunotherapy (PICI), and the Chan Zuckerberg Initiative.

## Disclosures/Competing Interests

C.J.Y. is the founder of and holds equity in DropPrint Genomics (now ImmunAI) and Survey Genomics. He serves as a Scientific Advisory Board member for and holds equity in Related Sciences and ImmunAI and is a consultant or has consulted for Maze Therapeutics, TReX Bio, HiBio, ImYoo, Kiragen, and Santa Ana Bio. Additionally, C.J.Y. is an Innovation Investigator at the Arc Institute. He has received research support from the Parker Institute for Cancer Immunotherapy, Chan Zuckerberg Initiative, Chan Zuckerberg Biohub, Genentech, BioLegend, ScaleBio, and Illumina.

