## Supplementary Table for "A genome-wide CRISPR activation map of surface protein expression in human CD4 T cells"

**Supplementary Table 1.** Protein assignments from the hierarchical clustering of NTC cells expression correlation.

=== Activation (n=42) ===

B7-H4, C5L2, CD101, CD107a, CD134, CD137, CD151, CD152, CD154, CD162, CD186, CD194, CD2, CD223, CD226, CD25, CD257, CD26, CD278, CD3, CD314, CD317, CD35, CD45R-B220, CD45RA, CD49d, CD52, CD55, CD63, CD70, CD71, CD73, CD81, CD82, CD83, CD86, CD98, HLA-DQ, HLA-DR, HLA-DR-DP-DQ, integrinB7, TCR-Va7.2

=== Adhesion (n=39) ===

CD10, CD102, CD105, CD11c, CD123, CD146, CD150, CD166, CD171, CD196, CD218a, CD273, CD274, CD31, CD319, CD321, CD33, CD36, CD43, CD45RB, CD49a, CD49b, CD5, CD58, CD62L, CD8, CD84, CD85j, CD9, CD90, CD96, CD99, EGFR, KLRG1, MERTK, MICA-MICB, Notch1, Notch2, TCR-Vd2

=== Cytokine/Chemokine (n=39) ===

CD104, CD115, CD119, CD120b, CD124, CD138, CD144, CD158, CD158b, CD158d, CD178, CD183, CD195, CD197, CD198, CD199, CD23, CD244, CD254, CD267, CD269, CD270, CD279, CD294, CD307c-FcRL3, CD324, CD335, CD336, CD37, CD74, CX3CR1, GARP, LAP, Mac-2 , MR1, TCR-Vb13.1, TCR-Vr9, TCRrd, TSLPR

=== Core (n=24) ===

anti-CD230-Prion, CD109, CD11a, CD18, CD184, CD200, CD224, CD29, CD352, CD38, CD4, CD44, CD45, CD45RO, CD46, CD47, CD48, CD49f, CD54, CD59, CD69, CD95, GP130, HLA-ABC

=== Innate-like (n=57) ===

CD103, CD106, CD117, CD11b, CD122, CD126, CD127, CD131, CD135, CD14, CD158e1, CD158f, CD16, CD161, CD185, CD192, CD193, CD1d, CD21, CD24, CD243, CD253, CD268, CD27, CD272, CD28, CD30, CD305, CD32, CD337, CD34, CD357, CD360, CD365, CD366, CD39, CD40, CD41, CD42b, CD56, CD57Recombinant, CD64, CD7, CD72, CD94, CLEC1B, FolateReceptorB, GPR56, HLA-A2, HLA-E, NKp80, Notch3, TCR-Va24-Ja18, TCRab, TIGIT, TIM-4, XCR1
