## Supplementary Figures for "A genome-wide CRISPR activation map of surface protein expression in human CD4 T cells"

**Supplementary Fig. 1. Gating strategy for SCITO-Perturb-seq**

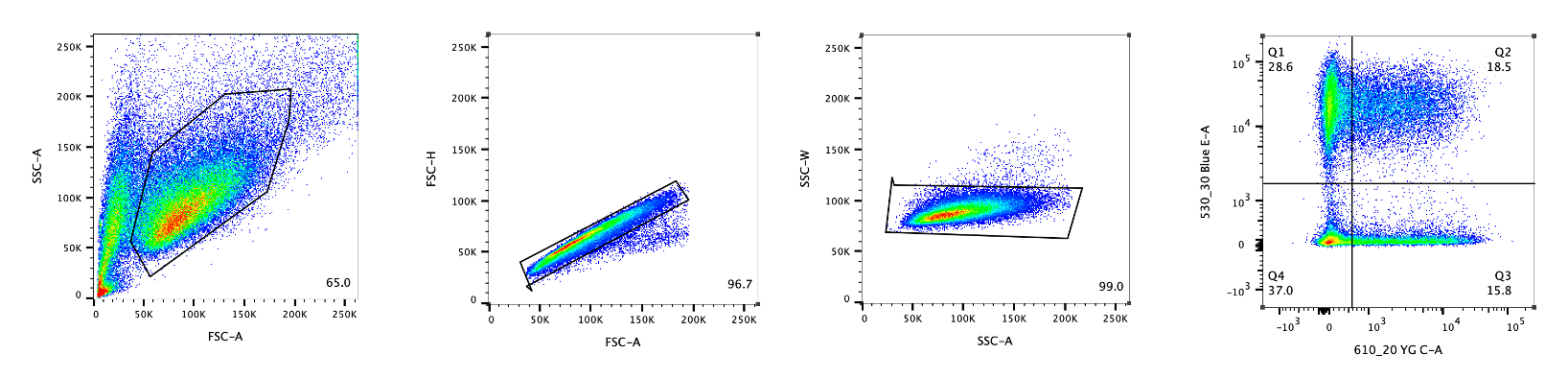

Representative sequential flow cytometry plots are shown from the left to right. Lymphocytes were gated for singlets and double-positive populations (YG-C channel for dCas9 vector and Blue E channel for the guide library) are sorted for library construction.

**Supplementary Fig. 2. Pilot experiment for scalable CRISPR screen**

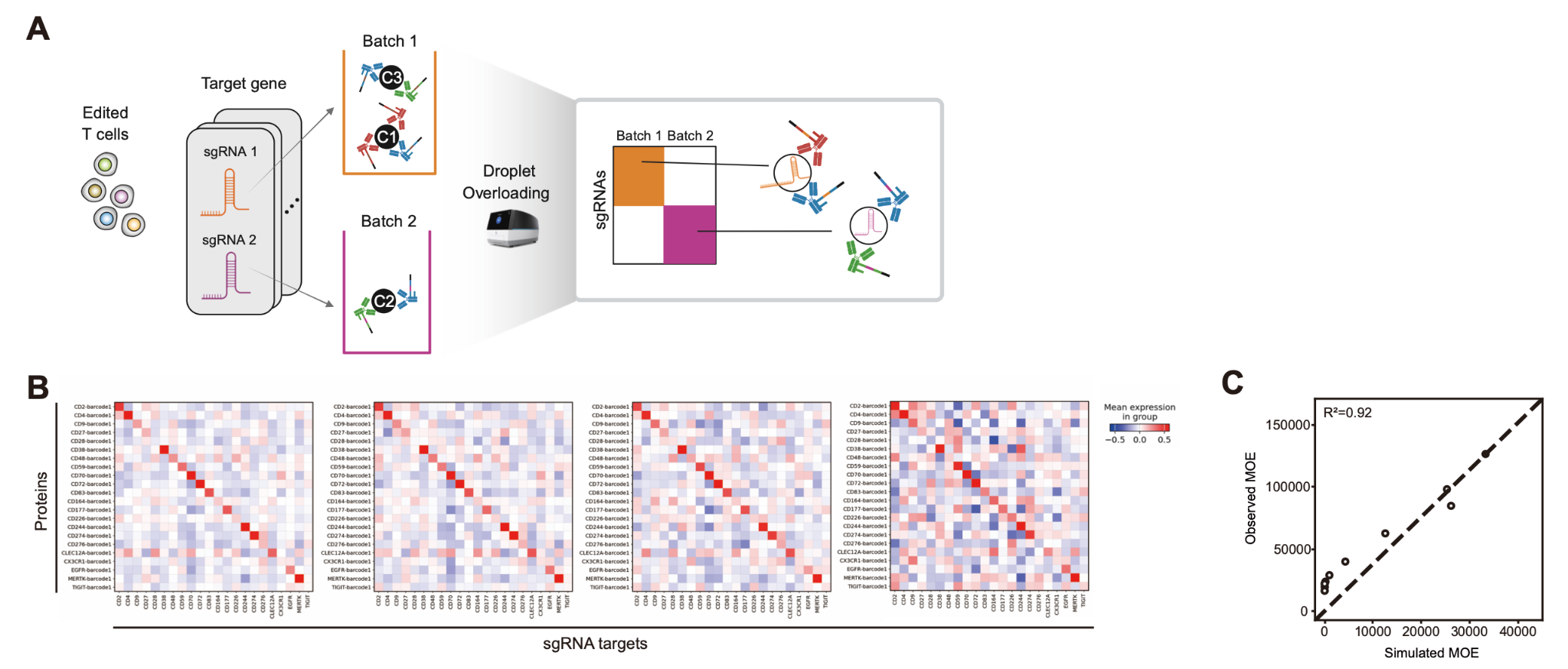

(A) Experimental design for positive control pilot experiment targeting 22 surface proteins.

(B) Heatmap of the average normalized expression of 22 perturbation target proteins in perturbed subpopulations defined by the sgRNA assignment. The average normalized expression is computed for all cells composed of ~790 cell subpopulations assigned per sgRNA(left) and for the cells downsampled by the 50%, 30%, and 10%(right) ratio, respectively.

(C) Scatter plot of simulated (x-axis) and observed (y-axis) multiplicity of encapsulation(MOE) averaged across 10 SCITO-Perturb-seq pools. Simulated MOE is computed assuming the Poisson loading of 500,000 cells on 200,000 droplets and repeated 10 times representing each pool.

**Supplementary Fig. 3. Direction and cell count of non-significant self-targeted pairs are consistent with limited power rather than absent effects.**

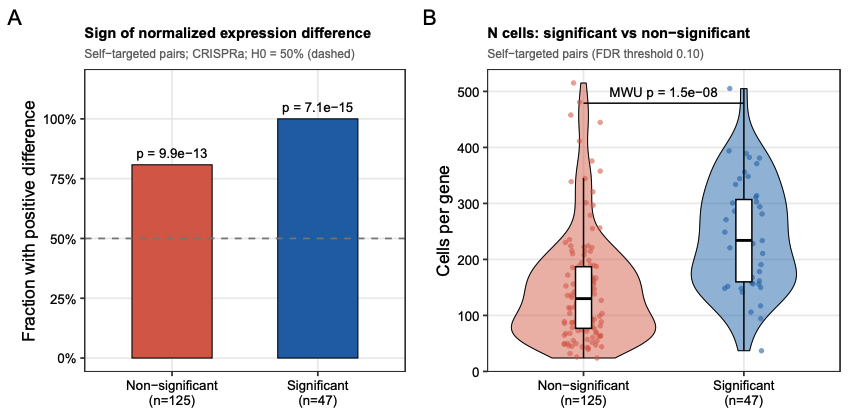

(A) Sign of the normalized expression difference for self-targeted gene-protein pairs, split by significance. Self-targeted pairs are classified as non-significant (per-protein BH-adjusted q ≥ 0.10, n = 125) or significant (q < 0.10, n = 47). Bars show the fraction of pairs with a positive difference; the dashed line marks the 50% null expectation. P values are from a one-sided binomial test (H₀ fraction positive = 0.5). (B) Cells per gene for the same significant and non-significant self-targeted pairs. Each point is one gene-protein pair, positioned by the total number of cells assigned to the gene across all its sgRNAs (genes with fewer than 20 cells were excluded from the analysis). Violins show the kernel density, boxes show median and interquartile range, and whiskers extend to 1.5 × IQR. The bracket shows a one-sided Mann-Whitney U test (H1 significant pairs have more cells; p = 1.5 × 10⁻8).

**Supplementary Fig. 4. Metrics for selecting factorization rank in the semi-NMF analysis**

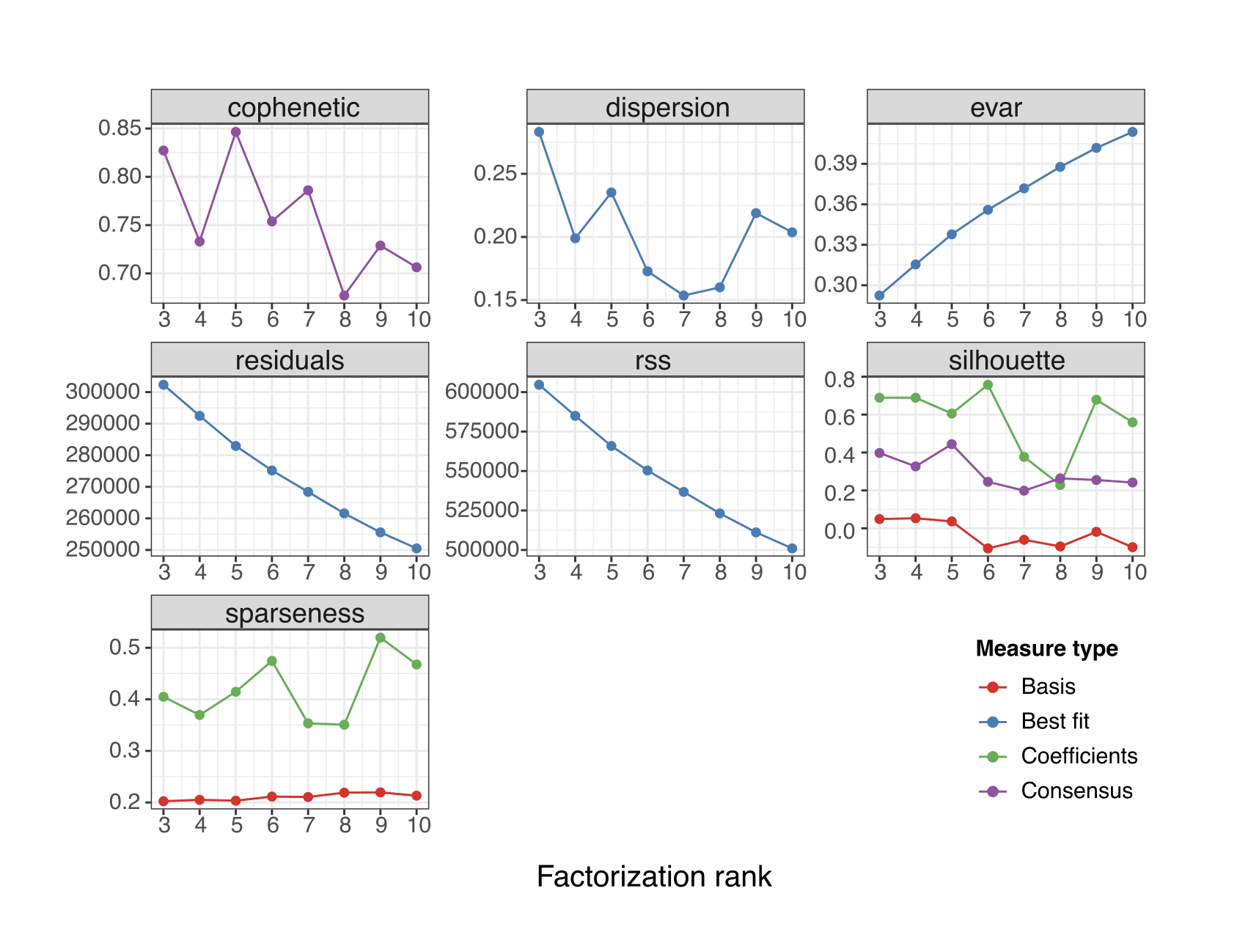

Multiple quality metrics were evaluated across factorization ranks (k = 3-10) to guide model selection. The cophenetic correlation summarizes how consistently samples cluster across repeated runs, with higher values indicating more reproducible sample assignments. Dispersion reflects how decisively samples are assigned in the consensus matrix, where values closer to 0 or 1 indicate clearer partitioning. The explained variance (evar) captures the proportion of total signal recovered by the factorization and increases steadily with rank. In contrast, residuals and residual sum of squares (RSS) quantify reconstruction error and decrease as additional components are introduced. These curves are therefore interpreted by identifying a point of diminishing returns. Silhouette scores provide a measure of cluster separation, with higher values indicating stronger within-cluster coherence relative to between-cluster similarity. Results are shown for basis (red), best fit (blue), coefficients (green), and consensus (purple). Finally, sparseness characterizes how concentrated the learned factors are, with higher values corresponding to more localized and interpretable components.

**Supplementary Fig. 5. Per-module stability of the semi-NMF decomposition under gene resampling.**

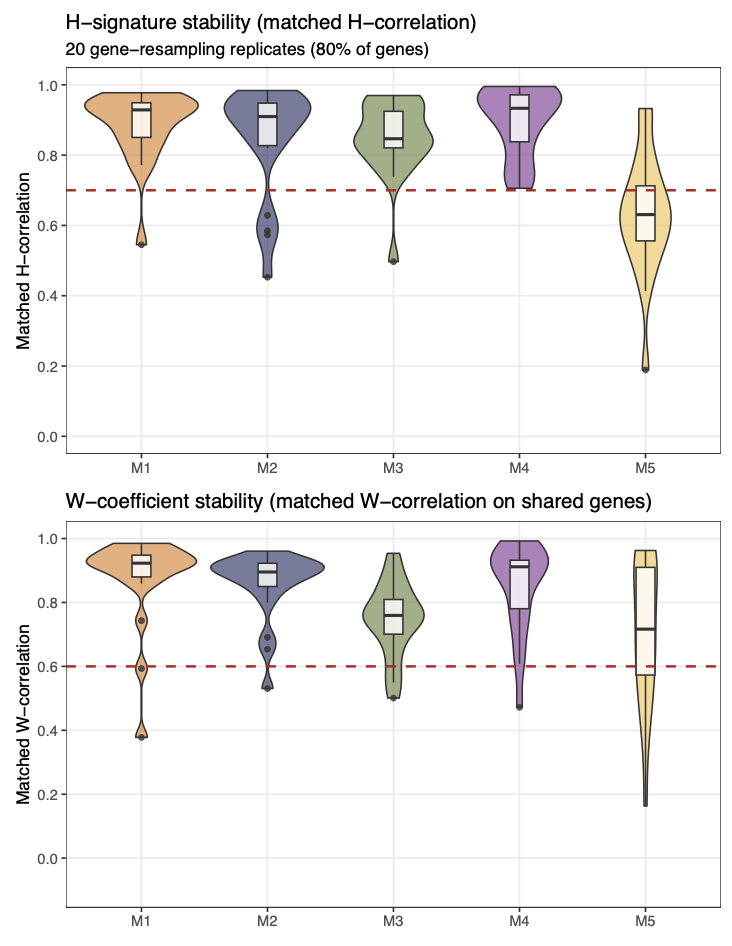

Twenty bootstrap replicates were generated by refitting semi-NMF on 80% gene subsets at K = 5, with modules aligned to the reference decomposition by Hungarian matching on H-signature correlation. Top, matched H-signature correlation per module. Bottom, matched W-coefficient correlation on shared genes per module. Boxes show median and interquartile range; dashed lines mark reference thresholds (0.70 for H, 0.60 for W). Modules M1 to M4 exceed the H threshold (medians 0.85 to 0.93); M5 falls below it (median 0.63), reflecting its low-amplitude protein signature.
